# Cotranscriptional RNA strand displacement underlies the regulatory function of the *E. coli thiB* TPP translational riboswitch

**DOI:** 10.1101/2022.08.24.505126

**Authors:** Katherine E. Berman, Russell Steans, Laura M. Hertz, Julius B. Lucks

**Affiliations:** Interdisciplinary Biological Sciences Graduate Program, Northwestern University, Evanston, IL 60208; Department of Molecular Biosciences, Northwestern University, Evanston, IL 60208; Department of Chemical and Biological Engineering, Northwestern University, IL, 60208; Center for Synthetic Biology, Northwestern University, Evanston, IL, 60208

**Keywords:** Riboswitches, strand displacement, RNA folding, translation regulation

## Abstract

Riboswitches are cis-regulatory RNA elements that regulate gene expression in response to ligand through the coordinated action of a ligand-binding aptamer domain (AD) and a downstream expression platform (EP). Previous studies of transcriptional riboswitches have uncovered diverse examples that utilize cotranscriptional strand displacement to mediate the switching mechanism. The coupling of transcription and translation in bacteria motivates the intriguing question as to whether translational riboswitches can utilize the same mechanistic features. Here we investigate this question by studying the *Escherichia coli thiB* thiamine pyrophosphate (TPP) riboswitch. Using cellular gene expression assays, we first confirmed that the riboswitch acts at the level of translational regulation. Deletion mutagenesis showed the importance of the AD-EP linker sequence for riboswitch function, which based on sequence complementarity with the AD P1 stem suggested the possibility of an intermediate structure reminiscent of transcriptional riboswitches that exploit strand displacement. Point mutation analysis of this intermediate structure, followed by designed changes to P1, supported a strand displacement mechanism for *E. coli thiB*. This work provides an important new example of diverse riboswitch AD-EP combinations that exploit this switching mechanism.

## INTRODUCTION

Riboswitches are RNA cis-regulatory elements that control gene expression in response to a ligand such as metabolites, ions, and other small molecules (Breaker 2012; McCown et al. 2017). Typically, riboswitches achieve their function through the coordinated action of two domains: the aptamer domain (AD) which folds into a structure that binds a particular ligand, and the expression platform (EP) which converts ligand binding into a gene regulatory outcome. With this architecture, riboswitches are classified by the cognate ligands they recognize, the aptamer structural motifs they use to do so, and the gene regulatory mechanisms employed by the EPs (Roth and Breaker 2009).

While much is known about how ADs bind ligands from structural and biophysical studies (Ray et al. 2018; Roth and Breaker 2009; Serganov and Patel 2012), much less is known about how ligand binding to the AD communicates through the EP to enact a regulatory decision (Garst et al. 2011). This is partly due to the diversity of EPs, with EPs known to be able to regulate diverse aspects of gene expression. For example, EPs can control transcriptional termination by forming a rho-independent terminator hairpin or exposing rho-termination factor binding site (Winkler 2003; Ray-Soni et al. 2016). Alternatively, EPs can regulate translation initiation through structures that occlude or expose a ribosome binding (RBS) (Nou and Kadner 2000), or RNA degradation by controlling exposure to RNase E sites (Caron et al. 2012; Winkler et al. 2002). Furthermore, the same AD can support the function of diverse EPs such as in thiamine pyrophosphate (TPP)-sensing riboswitches for which there are examples that regulate transcription (Sudarsan et al. 2005; Chauvier et al. 2017; Bastet et al. 2017), translation (Ontiveros-Palacios et al. 2007), and even mRNA splicing (Li and Breaker 2013; Cheah et al. 2007a).

Recent studies aimed at understanding riboswitch switching mechanisms have focused on transcriptional riboswitches, which necessarily must make their regulatory decision through forming a transcriptional terminator or anti-terminator before the RNA polymerase finishes transcribing the riboswitch (Watters et al. 2016; Hua et al. 2020). Because of this constraint, these riboswitches must fold in a cotranscriptional folding regime which has been shown to be important for proper riboswitch function (Chauvier et al. 2021; Hua et al. 2020; Perdrizet et al. 2012; Frieda and Block 2012; Scull et al. 2021). Cotranscriptional folding favors the formation of local RNA structures that can form in microsecond time scales as opposed to the milliseconds timescale of nucleotide incorporation by RNA polymerase (RNAP) (Ganser et al. 2019; Al-Hashimi and Walter 2008). However, structural rearrangements can occur during cotranscriptional RNA folding, which allows RNAs to escape kinetic traps and form more thermodynamically stable and sometimes functional folds (Pan and Woodson 1998; Pan et al. 1999; Pan and Sosnick 2006). One mechanism that allows RNAs to traverse free energy barriers to these rearrangements during the time window of transcription is strand displacement (Hong and Šulc 2019), a process by which an invading nucleic acid strand can base pair with a substrate strand by displacing a previously paired substrate:incumbent strand duplex (Hong and Šulc 2019). Strand displacement has been shown to underlie the ability of non-coding RNAs such as the *E.coli* Signal Recognition Particle to cotranscriptionally rearrange into long-range structures (Yu et al. 2021; Fukuda et al. 2020), as well provide a mechanism by which EPs can displace *apo-ADs* in diverse transcriptional riboswitches including those that sense ZTP (Strobel et al. 2019; Hua et al. 2020), fluoride (Watters et al. 2016), and guanine (Cheng et al. 2022). The formation of a ligand-bound *holo-AD* is then understood to block strand displacement, allowing transcriptional riboswitches to govern EP folding and gene regulation through AD-ligand interactions.

While strand displacement provides an efficient switching mechanism for transcriptional riboswitches, less clear is its importance for translational riboswitches (Scull et al. 2021). Previously, riboswitches which regulate gene expression with EPs that sequester the RBS have been shown to operate post-transcriptionally, in some cases performing their regulation through a ligand-mediated, thermodynamically-driven refolding event after transcription and before ribosome binding (Liberman et al. 2015; Polaski et al. 2016; Smith et al. 2010). Several comparative studies have also elucidated the similarities and differences between transcriptional and translational riboswitches that sense the same ligand, focusing on how ligand binding kinetics impact downstream gene regulation (Bhagdikar et al. 2020; Neupane et al. 2011; Suddala et al. 2013). For example, the translational versions of the *Vibrio vulnificus add* adenine riboswitch and the *Desulfurispirillum indicum metI* SAM riboswitch are both able to refold between the ligand bound and unbound state after transcription, as opposed to their transcriptional counterparts which may not refold after transcription (Bhagdikar et al. 2020; Neupane et al. 2011; Wickiser et al. 2005b, 2005a). While these are compelling examples, there is a potential for other mechanisms to exist for translational riboswitches. In particular, the coupling of transcription and translation in bacteria (Kohler et al. 2017) makes it possible that some translational riboswitches could operate in a cotranscriptional folding regime. This raises the intriguing question as to whether translational riboswitches could exploit some of the similar features of strand displacement switching mechanisms employed by known examples of transcriptional riboswitches (Bushhouse et al. 2022).

Here we aimed to investigate this question through studying the TPP riboswitch, the only known riboswitch class represented in all domains of life, with the second highest number of unique riboswitches identified (Moldovan et al. 2018; Cheah et al. 2007b; Li and Breaker 2013; Antunes et al. 2019), and diverse EPs that can control transcription, translation, degradation and splicing all with the same highly conserved AD (Bushhouse et al. 2022; Cheah et al. 2007a; Li and Breaker 2013). TPP is the active form of thiamine or vitamin B-1, produced by thiamine diphosphokinase, which catalyzes several biochemical reactions in the cell (Nakayama and Hayashi 1972; Jurgenson et al. 2009). The TPP AD is formed from a well-conserved three-way junction, consisting of a long-range P1 hairpin with two branched hairpins that comprise the ligand-binding pocket (Serganov et al. 2006) (Figure 1A, S1). The TPP molecule is oriented in the pocket in an extended conformation (Figure S1), with the thiamine and pyrophosphate moieties interacting with ligand binding sites with nucleotides in junction 2/3 and 4/5 respectively (Serganov et al. 2006). The P1 helix makes no direct contacts with TPP, but rather is stabilized by long-range ligand binding interactions between the two branched helices.

**Figure 1:**
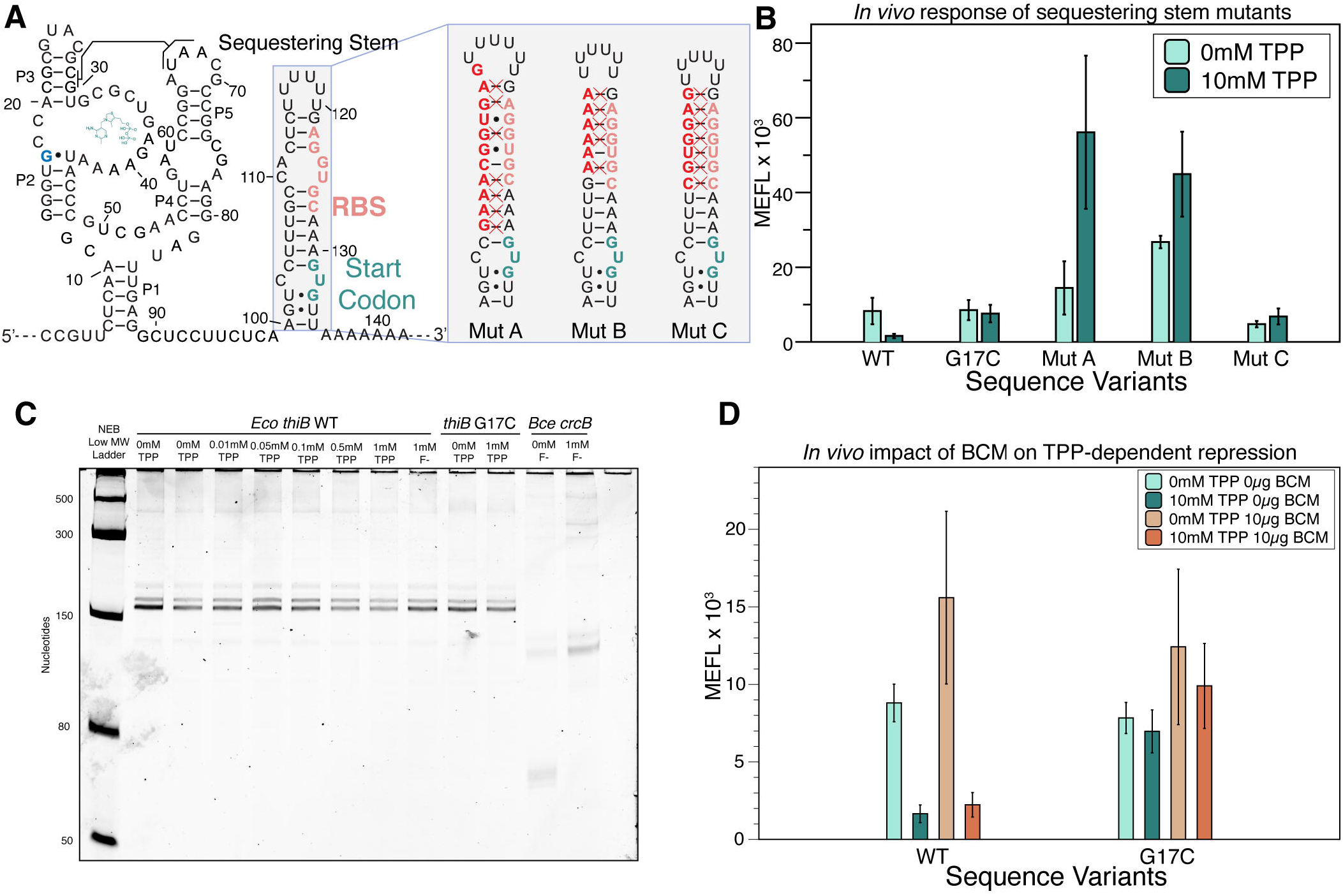
The *thiB E. coli* TPP riboswitch represses translation through a sequestering stem that includes the ribosome binding site. A) Proposed secondary structure of the ligand bound *E. coli thiB* TPP riboswitch based on the published crystal structure(Serganov et al. 2006), Rfam sequence alignment(Kalvari et al. 2017), and the sequestering stem secondary structure modeled using RNAstructure(Reuter and Mathews 2010). G17, which is known to participate in ligand binding is highlighted blue. The ribosome binding site (RBS) and the start codon for the naturally encoded downstream *thiB* gene are colored in red and green, respectively. The chemical structure of TPP is drawn next to the RNA, and a known tertiary interaction between the P2 and P3 stems is highlighted. Inset shows secondary structure models of three mutant sequestering stems designed to strengthen basepairing with the RBS. Mutated nucleotides are outlined in red with a red X indicating a base pair that is modeled to be disrupted by the mutation. Secondary structures determined by using RNAstructure version 6.4(Reuter and Mathews 2010). B) Wild type and mutant *thiB* riboswitch-regulated sfGFP expression in *E. coli* cells measured by flow cytometry. Units shown are molecules of equivalent fluorescein (MEFL) determined by flow cytometry. C) Single round *in vitro E. coli* RNA polymerase transcription assay measuring transcription products on a 10% Urea Polyacrylamide gel electrophoresis. The *Bce crcB* F-riboswitch, used as a positive control, terminates in the absence of F^-^ at an expected length of 82nt, and antiterminates in the presence of F^-^ at an expected length of 124 nt. The expected length of the full *thiB* transcription product is 173 nt. D) Wild type and mutant *thiB* riboswitch-regulated sfGFP expression in *E. coli* cells in the presence and absence of the rho-inhibitor bicyclomycin (BCM) measured by flow cytometry. Bar graphs in B and D represent mean values across three biological replicates, each performed in technical triplicate for nine total datapoints (n=9). Error bars represent the standard deviation from the mean. Data in C are n=2 representative gels (SI Figure S6, S9).

Currently, there are three identified *E. coli* TPP riboswitches which have been determined to use ribosome binding site (RBS) and start codon sequestration to down-regulate downstream operon gene expression, *thiM, thiC and thiB* (Winkler et al. 2002). However, recent studies have revealed dual control mechanisms in the *thiM* and *thiC* riboswitches that use NusG dependent RNA polymerase pausing, rho-dependent termination and RBS sequestration to perform their regulation (Chauvier et al. 2019, 2017; Bastet et al. 2017). While the *E. coli thiB* TPP riboswitch contains identified RNA polymerase pause sites and suggested rho-dependent termination in the downstream gene body, as well as sequence homology with *thiM* and *thiC*, the mechanism this riboswitch uses to downregulate gene expression is still unclear (Chauvier et al. 2017; Bastet et al. 2017). Intriguingly the *thiB* aptamer has been found to fold most efficiently in a cotranscriptional folding regime (Lang et al. 2007; Haller et al. 2013; Chauvier et al. 2021). We therefore sought to use the *E. coli thiB* TPP translational riboswitch as a model system to investigate the potential importance of cotranscriptional strand displacement for its folding mechanism.

Using cellular gene expression assays, we first confirmed that the *E. coli thiB* TPP riboswitch regulates at the level of translation in cells. Using a targeted mutagenesis approach, we next investigated deletions to the linker region connecting the AD and EP, and discovered a key sequence element in this region that was essential for function. Further exploration revealed that this sequence contained a complementary sequence to the P1 stem, indicating this sequence could base pair with the P1 stem to form alternative intermediate structures during the riboswitch folding pathway. This pattern of complementarity was reminiscent of similar patterns found in transcriptional riboswitches that were shown to utilize a strand displacement switching mechanism (Hong and Šulc 2019), which motivated detailed site-directed mutagenesis experiments with results consistent with a strand displacement mechanism underling *E. coli thiB* switching. Based on this mechanism, we designed sequence changes to the P1 stem to favor or disfavor the formation of the predicted intermediate structure facilitating strand displacement, and found that the function of the riboswitch changed to favor the ON or OFF state according to the direction of bias. Overall, the data support the conclusion that a cotranscriptional RNA strand displacement mechanism underlies the function of the *E. coli thiB* TPP riboswitch, providing an important new example of diverse riboswitch AD-EP combinations that exploit this switching mechanism.

## RESULTS

### The *Eco thiB* TPP riboswitch downregulates translation through a sequestering stem that occludes the ribosome binding site

We first sought to develop an assay to characterize *E. coli thiB* riboswitch gene regulation in response to TPP in *E. coli* cells. The *thiB* TPP riboswitch is predicted to function similarly to the *thiM* TPP riboswitch by occluding the ribosome binding site in the presence of ligand (Rodionov et al. 2002). A secondary structure model of the *thiB* riboswitch was generated from the consensus TPP riboswitch aptamer structure (Kalvari et al. 2017) and the predicted fold of the expression platform using RNAStructure (Reuter and Mathews 2010) (Figure 1A). Based on this knowledge, we designed and constructed a plasmid consisting of a constitutive *σ70 E. coli* RNA polymerase promoter followed by the *E. coli thiB* TPP riboswitch sequence and a super folder green fluorescent protein coding sequence (sfGFP) (Figure S2). The riboswitch coding sequence was designed to start at the transcription start site (Vogel et al. 2003) for the *thiBPQ* operon through the first 12 nucleotides of the *thiB* gene, with the downstream coding sequence included based on the predicted location of the sequestering stem that is thought to fold to occlude the RBS in the riboswitch OFF state (Figure 1A) (Rodionov et al. 2002).

We next used this expression construct to optimize an *E. coli* flow cytometry assay to characterize sfGFP fluorescence as regulated by the *thiB* TPP Riboswitch (Figure S2,S3). An important part of this assay is a ligand unresponsive control that should produce constitutive fluorescence independent of TPP. To generate this mutant, we used the crystal structure of the well-studied *E. coli thiM* TPP riboswitch (Serganov et al. 2006), along with mutagenesis data (Ontiveros-Palacios et al. 2007), to generate a series of point mutations predicted to interfere with ligand binding but not overall aptamer structure (Figure S4). Constructs were transformed in *E. coli* TG1 chemically competent cells, grown overnight in LB media, and then subcultured for 6 hours in thiamine hydrochloride deficient M9 media in the absence or presence of indicated concentrations of TPP. While the wild-type sequence showed repression in the presence of TPP, all mutants tested showed a broken ‘ON’ phenotype with fluorescence levels similar to or above the wildtype sequence, indicating that the WT repression was due to ligand interactions with the riboswitch and not TPP-dependent toxicity. We chose to use mutant G17C for additional experiments as it showed the most similar sfGFP expression to the wildtype *thiB* riboswitch with no ligand.

We next performed a dose response curve comparing the wildtype and G17C mutant riboswitch to determine the optimum concentration of TPP for downstream experiments, which revealed that 10mM TPP provided distinguishable repression between these two constructs (Figure S5). We next tested the importance of expression platform structural context on *thiB* repression. We hypothesized the formation of the predicted EP hairpin blocked ribosome binding to the ribosome binding site, therefore repressing gene expression. To test this hypothesis, we created a set of mutants that were designed to strengthen base pairing with the RBS in this sequestering stem (Figure 1A). Fluorescence characterization of these mutants showed that they resulted in higher fluorescence than the wildtype sequence, as well as no fluorescence repression due to TPP, indicating that base-pairing context with the RBS within the proposed sequestering stem is important for the *thiB* repression mechanism (Figure 1B).

To further confirm that *thiB* regulates at the level of translation, we tested whether this RBS sequestering stem could also function as an intrinsic transcriptional terminator by completing single-round *in vitro* transcription at increasing concentrations of TPP. In this assay, we observed only full-length transcription products with increased concentrations of TPP, indicating that the riboswitch does not function through transcriptional termination in these conditions (Figure 1C, Figure S6). Previous studies have shown *thiC* and *thiM E. coli* TPP riboswitches repress gene expression through dual control mechanisms that use both an RBS sequestering hairpin and rho-dependent transcriptional termination (Bastet et al. 2017; Chauvier et al. 2017). To test whether this could also be true for *thiB*, we tested whether the rho inhibitor, bicyclomycin (BCM) altered the riboswitch’s repression due to TPP. Characterization of the wildtype riboswitch sequence in cultures where BCM was added in the subculture M9 media at 25ug/mL, showed a similar repression in the presence of TPP, suggesting that rho is not important for riboswitch function (Figure 1D).

Taken together, these results demonstrates that the *thiB* TPP riboswitch regulates at the level of translation in our experimental context, and that base pairing interactions with the RBS within a putative sequestering stem are important for this regulation.

### Nucleotides in the linker region are essential for *thiB* gene regulation

Previous work has shown that sequences directly after the P1 helix can play a role in riboswitch mechanisms through the formation of intermediate structures that compete with EP folding to enact the switch (Cheng et al. 2022). Consequently, we next sought to determine whether this sequence could play a role in the *thiB* switching mechanism. We were specifically interested in investigating nucleotides 90-99, a non-conserved region that is predicted to form a single stranded ‘linker’ between the aptamer and the expression platform sequestering stem in the *thiB* OFF state (Figure 2A). While this ten-nucleotide sequence does not have sequence complementarity with nucleotides in the model of the sequestering stem, we sought to investigate whether this region, or portions of this region, are essential for *thiB* function. To address this question, we performed functional mutagenesis of this region by deleting or randomizing portions of the sequence (Figure 2B). Upon deletion or randomization of the entire 10 nucleotide sequence, we observed total loss of sfGFP expression in the presence or absence of TPP, indicating that the riboswitch was broken in the OFF state (Figure 2C). We next investigated deleting or mutating portions of the linker region. Deleting or randomizing the 5’ half (nucleotides 90-94) resulted in mutants that demonstrated repression in the presence of TPP similar to the wildtype sequence (Figure 2C), indicating that this sequence was not essential for riboswitch function. However, deletion or randomization of the 3’ half (nucleotides 95-99) showed the broken OFF phenotype, indicating that this region is essential for riboswitch function. Taken together, these results demonstrate that the linker plays an important role in *thiB* regulation.

**Figure 2:**
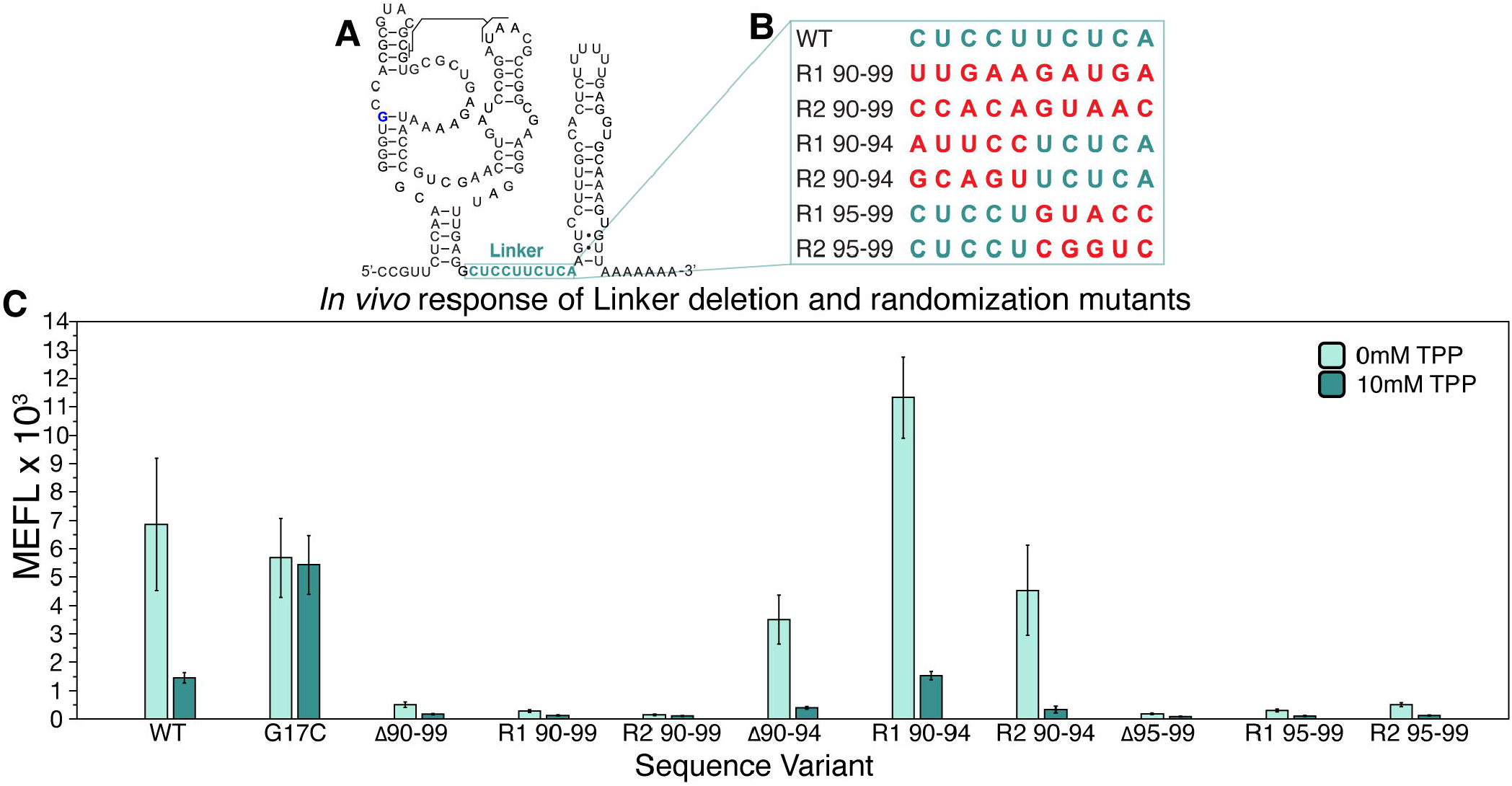
A specific portion of the linker region in the *E. coli thiB* TPP riboswitch is essential for TPP-dependent repression. A) Secondary structure model of the *thiB E. coli* TPP riboswitch after Figure 1A, with linker sequence (nts 90-99) outlined in light green. B) A table of sequence randomizations tested within the linker sequence. C) Flow cytometry assay of plasmid sfGFP expression regulated by the ThiB riboswitch sequence variants. X-axis labels indicate the riboswitch sequence variant tested (WT = wildtype, G17C = ligand unresponsive mutant, Δ = deletion). Bar graphs represent mean values across three biological replicates, each performed in technical triplicate for nine total datapoints (n=9). Error bars represent the standard deviation from the mean.

### Linker region mutations suggest strand displacement is important for *thiB* gene regulation

The above results demonstrated the importance of the 3’ half of the linker region for *thiB* function. A closer analysis of this sequence revealed that these nucleotides (95-99) are complementary to the 3’ side of P1 (nts 85-89), potentially able to form an intermediate ‘anti-sequestering’ hairpin together (Figure 3A). In addition, this linker region sequence is identical to the 5’ side of the P1 stem (nts 5-9), suggesting that the P1 helix and the anti-sequestering helix could be mutually exclusively basepaired regions within the riboswitch. Notably, such internal competing structures have been identified in other riboswitch mechanisms that appear to leverage strand displacement in their switching mechanisms (Strobel et al. 2019; Cheng et al. 2022).

**Figure 3:**
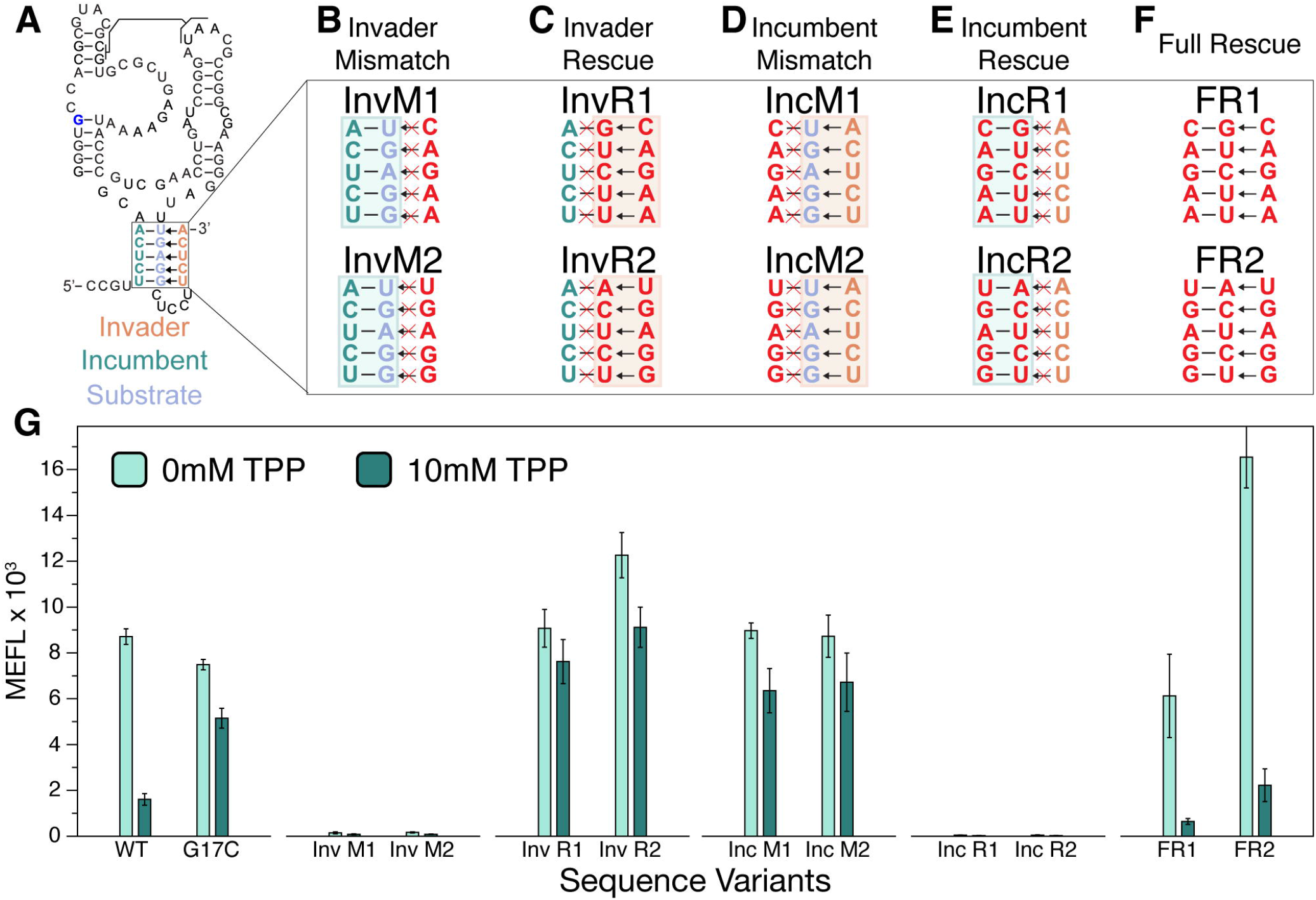
Complementary base pairs within P1 and within the anti-sequestering stem are required for *E. coli thiB* TPP riboswitch TPP-dependent repression. A) Secondary structure model of the *thiB E. coli* TPP riboswitch after Figure 1A including the proposed top portion of the anti-sequestering stem drawn as invading the P1 helix. Invader, incumbent and substrate strands are labeled according to the proposed strand displacement mechanism where the incumbent:substrate duplex forms the P1 stem of the aptamer and the incumbent:invader duplex forms the top of the anti-sequestering stem. Arrows indicate the proposed strand invasion mechanism where the invader displaces the incumbent during anti-sequestering stem folding. (B-F) Mutant nucleotides are depicted in red, while wildtype nucleotides are green (incumbent), orange (invader), and blue (substrate). Mutations were designed to break the invasion of the anti-sequestering stem (B, invader mismatch, InvM), rescue the invasion and break P1 (C, invader rescue, InvR), break P1 and allow invasion by the anti-sequestering stem (D, incumbent mismatch, IncM), rescue P1 and break the invasion (E, incumbent rescue, IncR), and rescue all base pairing interactions with sequences different than the wild-type sequence (F, full rescue, FR). G) Flow cytometry assay of plasmid sfGFP expression regulated by the ThiB riboswitch sequence variants. X-axis labels indicate the riboswitch sequence variant tested (WT = wildtype, G17C = ligand unresponsive mutant). Bar graphs represent mean values across three biological replicates, each performed in technical triplicate for nine total datapoints (n=9). Error bars represent the standard deviation from the mean.

Strand displacement is a process in which nucleotide strands basepair at the cost of displacing another previously bound nucleotide strand (Hong and Šulc 2019). Specifically an internal helix, composed of a ‘incumbent’ strand:’substrate’ strand duplex, can be broken apart when an ‘invader’ strand forms the competing invader:substrate duplex. Within the *thiB* system, this nomenclature can be used to label the 5’ side of the P1 (nts 5-9) as the incumbent strand, the 3’ side of P1 (nts 85-89) as the substrate strand, and nts 95-99 of the linker region as the invading strand (Figure 3A). Using this framework, we hypothesized that mutations to these strands would bias riboswitch function according to which structure they favored: mutations that favored incumbent:substrate pairing would favor the formation of P1 and the sequestering stem and bias riboswitch function to the OFF state, while mutations that favored substrate:invader pairing would favor the formation of the anti-sequestering stem and favor the riboswitch ON state.

To test these hypotheses, we constructed sets of mismatch mutants designed to break and rescue specific substrate:incumbent and substrate:invader base pairs and characterized their gene expression. As predicted, when the incumbent:substrate interaction was favored, the riboswitch function was broken in the OFF state (Figure 3B, E, G). In contrast, when the substrate:invader interaction was favored, the riboswitch was broken in the ON state (Figure 3C,D,G). Finally, a full rescue of both incumbent:substrate and substrate:invader interactions, such that the mutually exclusive hairpin structures are both possible, fully rescued the ability of the thiB sequence to regulate expression in response to TPP, although with varying dynamic range than the wildtype sequence (Figure 3F,G).

Taken together, these results support the importance of pairing between the 5’ and 3’ sides of P1 and the 3’ side of P1 with the linker to form an anti-sequestering hairpin for riboswitch switching. These results match those seen in other riboswitch systems thought to utilize strand displacement to form intermediate structures in their switching mechanism (Cheng et al. 2022), strongly suggesting a similar strand displacement mechanism is present in the *E. coli thiB* mechanism.

### Lengthening or shortening the P1 stem or the anti-sequestering stem bias TPP riboswitch regulation

The above results demonstrate the importance of overlapping sequence complementarity between P1 and the anti-sequestering stem for proper riboswitch function. Based on this data, we predicted that manipulating the length of either helix could impact strand displacement kinetics and therefore riboswitch function. Specifically, we hypothesized that strengthening P1 by adding base pairs, or shortening the anti-sequestering stem with mismatches, could inhibit anti-sequestering hairpin formation and bias the riboswitch to the OFF state even in the presence of ligand. Conversely, promoting the formation of the anti-sequestering stem – either through P1 weakening mutations or anti-sequestering stem strengthening mutations – could bias the riboswitch in the ON state.

To test these hypotheses, we lengthened the P1 stem by adding complementary nucleotides to the 5’ side which could base pair with the loop of the anti-sequestering stem (Figure 4A). These mutants expressed low amount of sfGFP both with and without TPP, confirming that strengthening the P1 helix with more basepairs favored the OFF pathway (Figure 4A). We next shortened the P1 helix by making 5’ side mismatch mutations (Figure 4A), which resulted in mutants that repressed in the presence of TPP, but with overall higher sfGFP expression both in the presence and absence of TPP. To demonstrate this effect was independent of sequence, we designed a second set of P1 shortening mutants, which had the same impact on sfGFP expression and repression due to TPP (Figure S7). This further supported our hypothesis that weakening the P1 stem biases the folding pathway towards the ON state (Figure 4A). These results demonstrate that the P1 helix length has an inverse relationship *thiB* TPP riboswitch overall expression.

**Figure 4:**
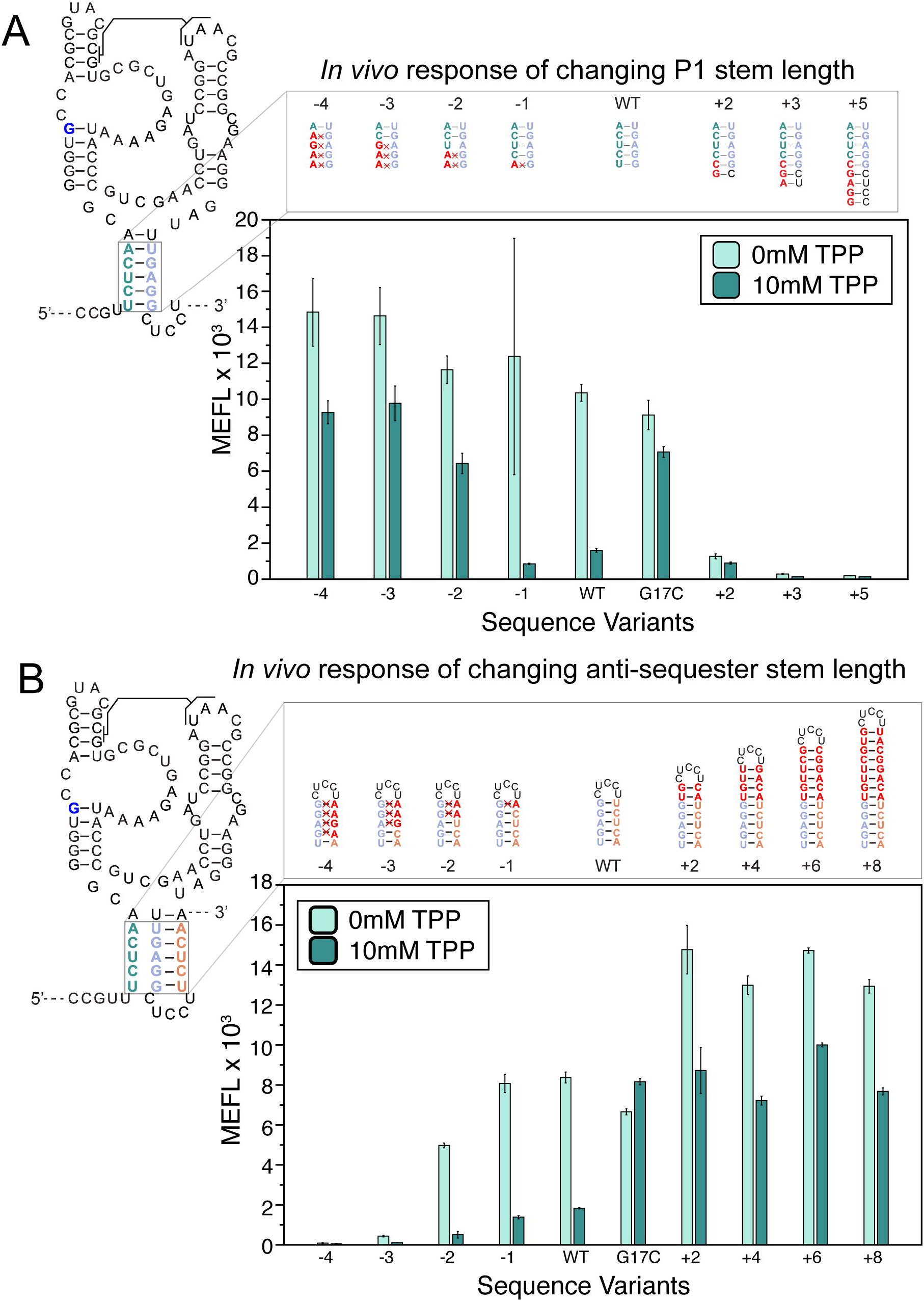
Changing the length of P1 and the anti-sequestering stem have the opposite effects on *E. coli thiB* TPP riboswitch function. A) P1 stem mutants that lengthen and shorten the P1 stem. Lengthening mutants change nucleotides in the leader sequence before the riboswitch to extend P1. Mismatches on the 5’ end shorten the P1 helix from the bottom. Secondary structure depictions of the mutations are depicted above flow cytometry assay data of plasmid sfGFP expression regulated by the *thiB* riboswitch P1 sequence variants. X-axis labels indicate the riboswitch sequence variant tested (WT = wildtype, G17C = ligand unresponsive mutant). B) Sequence mutants for the anti-sequestering stem. Mutants to the left of the wildtype hairpin are mismatch mutations which shorten the anti-sequestering helix, while mutants to the right extend the helix. Secondary structure depictions and flow cytometry assay data as in (A). X-axis labels indicate the riboswitch sequence variant tested (WT = wildtype, G17C = ligand unresponsive mutant). Bar graphs represent mean values across one biological replicate, each performed in technical triplicate for three total datapoints (n=3). Error bars represent the standard deviation from the mean.

Next, we designed mutations to weaken and strengthen the anti-sequestering stem, testing our hypothesis that anti-sequestering stem length has a direct relationship with *thiB* TPP riboswitch expression (Figure 4B). As predicted, mismatch mutations to the anti-sequestering stem 3’ side which shortens the anti-sequestering stem by three or more base pairs result in always OFF phenotypes where sfGFP expression is low both in the presence and absence of TPP. We next designed a set of mutants which added basepairs to the top of the anti-sequestering helix two basepairs at a time. As predicted, these anti-sequestering stem lengthening mutants resulted in higher overall sfGFP expression than the wildtype riboswitch. However, interestingly these anti-sequestering stem lengthening mutants were able to repress in the presence of TPP and increasing the length of the anti-sequestering stem beyond four basepairs did not increase the sfGFP expression above the two basepair addition. This suggests there is a limit to the impact anti-sequestering stem lengthening may have on riboswitch expression. Similar to our previous results, we created a second set of anti-sequestering stem lengthening and shortening mutants to test overall sequence bias, and the second set of mutants had similar results (Figure S8). Overall, these results matched the behavior of the corresponding P1 mutations, demonstrating the direct relationship between anti-sequestering stem length and *thiB* TPP riboswitch expression.

Taken together, these results show that the riboswitch regulatory decision can be biased in opposite directions through mutations that favor the formation of P1 or the anti-sequestering stem, pointing to the central role these structures play in determining riboswitch fate and providing further evidence of a strand displacement mechanism.

## DISCUSSION

In this work, we present evidence that the *E. coli thiB* TPP riboswitch regulates gene expression via a translational regulatory mechanism, but switches functional conformations using a strand displacement mechanism. Using cellular gene expression assays, we were able to first verify that *thiB* acts at the level of translational regulation (Figure 1). Mutagenesis of the *thiB* linker region that connects the aptamer domain to the expression platform revealed a sequence element that when deleted resulted in riboswitches that were broken OFF, indicating it’s important for the switching mechanism (Figure 2). The complementarity of this sequence element to the 3’ side of the aptamer P1 helix led to the hypothesis of an intermediate ‘anti-sequestering’ stem that competes with P1 formation to enact the regulatory decision. Characterization of riboswitch mutants that were designed to bias the formation of P1 or this anti-sequestering stem, either through point mutations (Figure 3) or lengthening/shortening mutations (Figure 4), supported a model of a strand displacement mechanism for the *thiB* TPP riboswitch mechanism. Overall, these results support a model of strand displacement as the method of switching between two alternative EP folds within the *thiB* TPP riboswitch.

This work represents another example of strand displacement being important for riboswitch mechanisms, this time in the context of a translational riboswitch. Interestingly, our results match some of those observed for transcriptional riboswitches such as the *B. subtilis yxjA* purine riboswitch, which was found to nascently form a central helix responsible for strand displacing its P1 helix of the aptamer in the absence of ligand while also outcompeting a second strand displacement process with the EP terminator hairpin (Cheng et al. 2022). Similarly, our mutational analysis suggests that the *thiB* anti-sequestering stem can strand displace the P1 helix in the absence of ligand to prevent the full formation of a functional RBS sequestering stem. These results are consistent with results from a recent cotranscriptional smFRET study, which found transcription to be essential for the *thiB* riboswitch to sense ligand, and that anti-sequestering hairpin formation defines the end of the ligand sensing transcription window (Chauvier et al. 2021).

While our functional mutagenesis supported an overall strand displacement mechanism that governs EP folding, our study design precluded us from investigating how TPP binding to the *thiB* AD blocks strand displacement of P1 to promote the formation of the sequestering stem. Other studies of riboswitches have demonstrated the importance of P1 hairpin length on ligand binding, sensing, and switching (Nozinovic et al. 2014; Drogalis and Batey 2020). Previous smFRET studies of the *thiM* TPP riboswitch have suggested distal binding of TPP in the binding pocket had a stabilizing effect on P1 by reducing residual dynamics of the helix (Haller et al. 2013). This same study found lengthening the P1 helix by 2 GC basepairs stabilized P3/L5 interactions, which could potentially contribute to understanding how TPP binding prevents strand displacement (Haller et al. 2013). While not tested here, this leads to a possible second event that could tune riboswitch function – as the anti-sequester stem competes with the P1 hairpin, the sequestering stem can also begin to nascently form downstream, creating a competition between anti-sequestering and sequestering stem formation that could ultimately tune riboswitch function as has been observed in the yxjA transcriptional riboswitch (Cheng et al. 2022).

This work could also shed light onto the evolution of riboswitch expression platform sequence. While aptamer sequences are highly conserved, expression platform sequences are known to show a high degree of variability (Moldovan et al. 2018). As seen in this work, changes to expression platform sequence can bias the riboswitch towards the ON or OFF state, indicating that expression platform sequence variation could be an evolutionary mechanism to tune riboswitch function. Studies that have uncovered the sequence determinants of efficient strand displacement (Hong and Šulc 2019) may shed light on this tuning as well and offer new synthetic routes to tuning riboswitch function.

Overall, this work starts to connect our understanding of the folding mechanisms of transcriptional and translational riboswitches showing, in the case of the *E. coli thiB* TPP riboswitch, that similar strand displacement mechanisms can be used by diverse riboswitches. As such, this study provides inspiration for further investigation of whether strand displacement is important in other translational riboswitch systems, and potentially through other RNA regulatory mechanisms(Bushhouse et al. 2022).

## MATERIALS AND METHODS

### Strains, Primers and Media

All plasmids were derived from the p15a plasmid backbone with chloramphenicol resistance. Plasmid sequences are listed in Supplementary Data File S1, and were created either using Gibson Assembly or inverse PCR (iPCR). Some strains were deposited in Addgene with ascension numbers listed in Supplementary Data File S1. All strains were grown in Difco LB broth for cloning and purification. Gibson assembly was used to add the *E.coli thiB* wt sequence downstream of the *E. coli* sigma 70 consensus promoter (annotated as J23119 in the registry of standard parts, Supplementary Data Table S1) followed by the super folder green fluorescent protein (sfGFP) sequence. In order to create different mutants using iPCR, primers designed to add mutant sequences were ordered from Integrated DNA Technologies (IDT). Then, 200μL PCR reactions were mixed with 1-10ng/μL of template DNA, 200μM dNTPs, 1X Phusion Buffer, 100nM each primer and 0.25μL of Phusion Polymerase (2000U/mL; NEB). PCR products were run on a 1% agarose gel to confirm desired length. Dpn1 (NEB) was used to digest template DNA and PCR clean up (Qiagen) used to purify PCR products. Digested and purified PCR products were then phosphorylated and ligated simultaneously using T4 PNK Enzyme (NEB), T4 DNA Ligase (NEB), and 10X T4 DNA Ligase buffer by incubating at room temperature for one hour. Ligation products were then transformed into NEBTurbo competent cells and plated on chloramphenicol LB agar plates and incubated at 37°C overnight. The next day, isolated colonies were inoculated in 4mL LB media cultures, miniprepped and then sequence confirmed using Sanger Sequencing (Quintara Biosciences).

### Flow Cytometry Data Collection of sfGFP Fluorescence Regulated by Riboswitches

For each experiment, the indicated constructs were transformed into *E. coli* strain TG1 (F’ [traD36 proAB lacIqZ ΔM15] supE thi-1 Δ(lac-proAB) Δ(mcrB-hsdSM)5(rK - mK -). Plasmids were transformed into chemically competent *E. coli* cells, plated on Difco LB agar plates containing 34 ug/mL chloramphenicol, and incubated at 37°C for 16-18 hours. Plates were then placed at room temperature for 6-8 hours. Three colonies per plasmid were inoculated in 200uL of LB chloramphenicol (34 ug/mL) and incubated overnight at 37°C. Following this, subcultures were created by adding 4uL of the overnight culture to 196uL of pre-warmed M9 minimal media lacking thiamine hydrochloride (M9 Salts, 0.4% glycerol, 0.2% casamino acids, 2mM MgSO_4_, 0.1mM CaCl_2_) with the indicated growth conditions. TPP at the indicated concentration was added to pre-warmed M9 media from solid powder directly before the experiment. Each culture block was covered with a Breath-Easier sealing membrane (Sigma-Aldrich) and incubated at 37°C while shaking at 1000rpm (Vortemp Shaking Incubator) for 6 hours. After 6-hours of subculture incubation, cells were diluted 1:100 in Phosphate Buffered Saline (PBS) and kanamycin (50μg/mL). A BD Accuri C6 Plus flow cytometer fitted with a high-throughput sampler was then used to measure the sfGFP Fluorescence for each sample. *E. coli* events were collected using the FSC-A threshold set to 2000. sfGFP fluorescence was measured by collecting FITC signal from the FL1-A channel. Fifty thousand events were captured for each sample.

### Flow Cytometry Data Analysis

Flow cytometry data analysis was performed using FlowJo (v10.4.1). Cells were density gated using an ellipsoid gate around the FSC-A vs. SSC-A plot, and the same gate was used for all samples prior to calculating the geometric mean of fluorescence in the FITC-A channel. Arbitrary fluorescent values from the flow cytometer were converted to molecules of equivalent fluorescein (MEFL) by measuring the fluorescence from Spherotech rainbow 8-peak calibration beads (Cat# RCP305A). A calibration curve was then created by calculating the linear regression between the arbitrary relative fluorescent values from the flow cytometer and the known MEFL provided by the manufacturer. The geometric mean of the FITC channel peaks after gating for *E. coli* cells was converted to MEFL by multiplying by the slope of the calibration curve and adding the y-intercept. Supplementary Figure 3 contains representative flow cytometry data from the measurement and calibration procedure.

### Single-round in-vitro transcription of linear templates with *E. coli* RNA Polymerase

Linear double-stranded DNA templates were prepared by 500uL PCR reactions using 50uL 10X Thermo Pol buffer (NEB), 10uL dNTP (10mM) (NEB), 2.5uL KEB.E74 (100uM) primer and KEB.E95 (100uM) primer, 2.5uL mini-prepped plasmid, and 2.5uL of Taq DNA Polymerase. The PCR reactions were then thermocycled at 95°C for 3 min, then 24 cycles of 95°C for 30 sec, 57°C for 1 min, and 68°C for 1 min and 30 seconds, and final extension at 68°C for 5 minutes. The PCR products were purified by ethanol precipitation and 1% agarose gel purified using the QIAquick Gel Extraction Kit (Qiagen). Single-round *E. coli* RNA polymerase *in vitro* transcription was then performed by mixing reactions on ice consisting of 100nM DNA template, 20mM Tris pH 8.0, 1uM EDTA pH 8.0, 1mM DTT, 50mM KCl, 1mM each NTP, 0.2ug BSA, 2 units of *E. coli* RNA Polymerase Holoenzyme (NEB), and the indicated concentration of ligand. We found that above 1mM TPP was toxic to *in-vitro* transcription. Each reaction was then incubated at 37°C for 10 minutes, then transcription initiated by adding 2.5uL of 100mM MgCl2 and 0.1mg/mL Rifampicin. After 30 seconds, the reaction was stopped with 75uL of TRIzol RNA Isolation Reagents (ThermoFisher Scientific). TRIzol extraction was then completed as directed to purify nucleic acids from the reaction. Extracted nucleic acids were resuspended in 43uL sterile, RNase free water, 5uL Turbo DNase 10x Buffer, and 2uL of Turbo DNase. Samples were then incubated at 37°C for 60 minutes and then RNA was purified with Trizol extraction as described previously. Purified RNA was resuspended in 10uL of sterile, RNase free water and then mixed with 10uL of RNA loading dye (7M Urea, 0.01% xylene cyanol and bromophenol blue). Then, samples and ssRNA LR ladder (NEB) were run on a 10% Urea PAGE at 16W for 90 minutes. The resulting PAGE was stained with SYBR Gold (Thermofisher Scientific) for 5 minutes and imaged using a ChemiDoc Imaging System (BioRad). Raw PAGE images are shown in SI Fig S9. Template sequences and primers used to create templates are listed in SI Table S1.

## AUTHOR CONTRIBUTIONS

K.E.B. and J.B.L conceived the projects. The data was curated, analyzed and checked by K.E.B and J.B.L.. K.E.B. and J.B.L. devised the methods as well as administered and validated the project. Flow cytometry experiments were completed by K.E.B and R.S.S. Single round *in vitro* transcription was completed by K.E.B. and L.M.H. Manuscript was written and edited by K.E.B. and J.B.L. Drafts were revised and edited by K.E.B., L.M.H., R.S.S., and J.B.L.

## ACKNOWLEDGEMENTS

We thank Adam Silverman and Luyi Cheng for experimental contributions to this study not included in this manuscript; Angela Yu for many helpful conversations about cotranscriptional RNA folding and strand displacement; and Alexis Reyes for careful feedback on manuscript drafts and figures.

